# Comment on ‘Valid molecular dynamics simulations of human hemoglobin require a surprisingly large box size’

**DOI:** 10.1101/563064

**Authors:** Vytautas Gapsys, Bert L. de Groot

## Abstract

A recent molecular dynamics investigation into the stability of hemoglobin concluded that the unliganded protein is only stable in the T state when a solvent box is used in the simulations that is ten times larger than what is usually employed. Here, we express three main concerns about that study. In addition, we show that with an order of magnitude more statistics, the reported box size dependence is not reproducible. Overall, no significant effects on the kinetics or thermodynamics of conformational transitions were observed.

## Introduction

A surprising effect of the simulation box size was recently reported for spontaneous hemoglobin transitions from the T to the R state in molecular dynamics simulations of deoxy hemoglobin (1). It was reported that the T state was stable for a period of 1 microsecond only in a relatively large simulation box with a cube edge length of 15 nm, whereas in smaller simulation boxes spontaneous transitions to the R state were observed on this timescale. Based on this, thermodynamic stabilization of the T state in the larger box was concluded, attributed to a change in the hydrophobic effect due to a change in the box size.

Here, we express three main concerns about the presented analyses. In addition, we present a systematic study of box size dependence in a variety of systems including hemoglobin, in which we observe no significant effect of the simulation box size on the kinetics or thermodynamics of conformational transitions.

## Results

### Kinetics rather than thermodynamics

First, we argue that the readout of transition times from T to R as applied is insufficient to allow for conclusions on the thermodynamic stability of the T state. According to rate theory, the transition times are primarily related to the activation barrier and the attempt frequency of the transition. Therefore, longer transition times could either be caused by a lowering of the attempt frequency or an increased barrier. In turn, an increased barrier (e.g. to leave the T state) can be achieved by a higher thermodynamic stability (lowering the well of the T state) but equally well by solely increasing the barrier and thus leaving the thermodynamic equlibrium between T and R unaltered. Without addressing the barrier and the attempt frequency, transition times to leave a state are thus insufficient to draw conclusions about the thermodynamic stability of that state. We thus argue that, based on the presented transition times, at most conclusions about the kinetics of the transition, but not on the thermodynamic stability of the T state can be drawn.

### Statistics

Second, for each of the four studied box sizes, a single MD simulation was carried out and a transition (or lack thereof) was reported. Based on the single data points, an estimate of the uncertainty in the reported transition times is lacking, impairing an assessment of the statistical significance of the observed differences. The statistical significance of the concluded box size dependence has thus not been shown. In redoing the simulations presented by El Hage et al. (1) using multiple replicas (20 for the 9 and 15 nm setup, 10 for the 12 nm setup, we find that there is considerable scatter in the transition times obtained in single MD simulations (see Fig 1). For example, we find that in the 9 nm box a substantial fraction does not undergo the transition within 1 *µ*s, rendering the absence of a transition in the single simulation of the 15 nm box in the El Hage et al. (1) study statistically insignificant. This is further underscored by the fact that out of 20 simulations we set up for the 15 nm system, 11 made the transition within 400 ns (Fig 1C). Consistent with this result, we find that the distributions of RMSD values to the T and R state crystal structures monitored at 400 ns and 1 *µ*s of simulation time are highly similar for the studied box sizes (Fig 1A,B).

**Figure 1:**
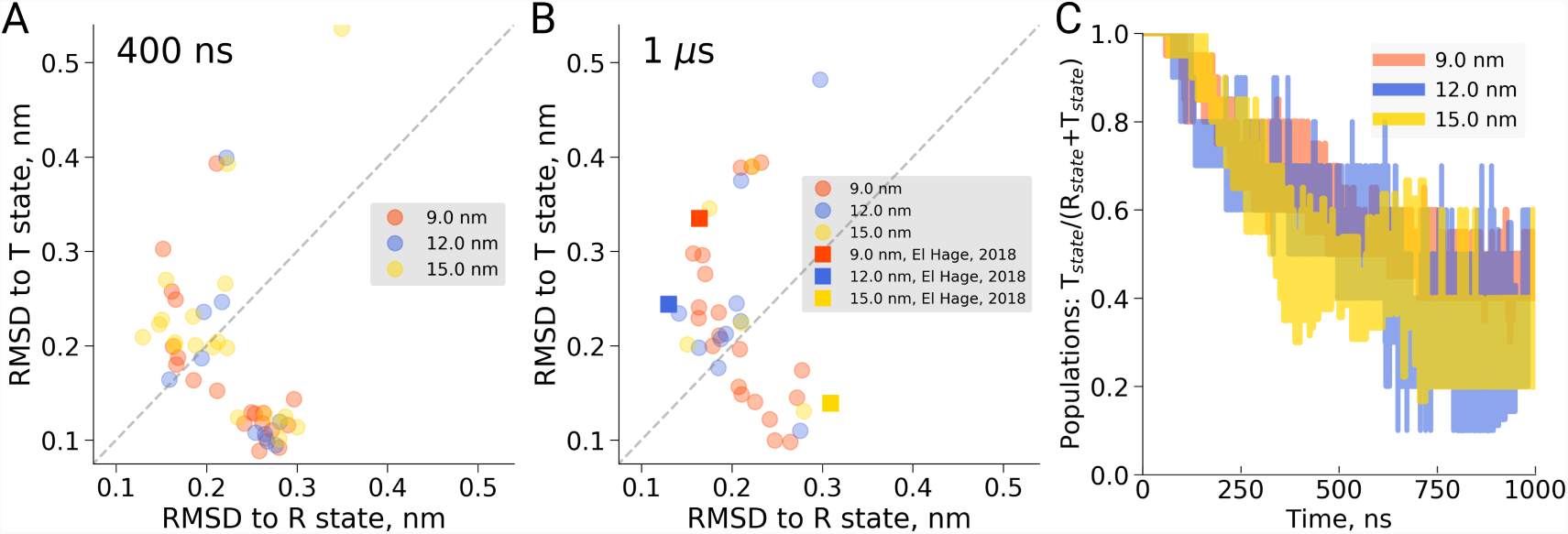
Hemoglobin simulation analysis. (A) RMSD of the simulation snapshots after 400 ns to the crystallographic T (2dn2) and R (2dn3) states. (B) RMSD of the simulation snapshots after 1 µs. (C) Fraction of trajectories remaining in T for simulations in the boxes of different size. Simulations were counted to have left the T state if more than half of the distance from the T state had been covered, as projected onto the difference vector between the T and R crystallographic states. For the 15 nm boxes, only 5 out of 20 simulations reached 1 µs, the others reached 400-750 ns.

### Hydrophobic effect

Third, the reported box size dependence was attributed to the hydrophobic effect, and analysed using solvent-solvent hydrogen bonds, solvent radial distribution function and water diffusion constants (1). Here, we are concerned that the presented analyses report not only on a possible hydrophobic or hydrodynamic effect, but also on a dilution effect caused by the different ratio of protein to water in differently sized boxes. Indeed, if we reanalyze the solvent radial distribution function (Fig 2A) or water-water hydrogen bonds (Fig 2B) and normalize based on the analysis volume (which we keep constant for the differently sized boxes) rather than by the box volume as was done originally, we find no significant box size effect. A similar dilution effect is observed for the diffusion constant. Averaging over the complete box represents a weighted average over restricted and bulk waters. For larger boxes, more water molecules are unrestricted by the protein and therefore behave bulk-like, yielding a larger weight of bulk water to the average. To disentangle this dilution averaging effect from possible box size effects, we estimated average diffusion constants for the 12 and 15 nm boxes from that of the 9 nm box, reweighting solely with the additional number of water molecules in the larger boxes and applying the bulk value of 5.95 10^5^*cm*^2^/s for those. As can be seen in Fig 2C, the resulting diffusion constants obtained this way are remarkably close to the ones originally computed for the 12 and 15 nm sized boxes, indicating that the observed diffences in average diffusion constant can be largely explained by the dilution effect and thus do not reflect a box size effect. This is consistent with earlier results from Yeh and Hummer (2) who showed that the box size dependence on water diffusion is small for box sizes larger than 4 nm.

**Figure 2:**
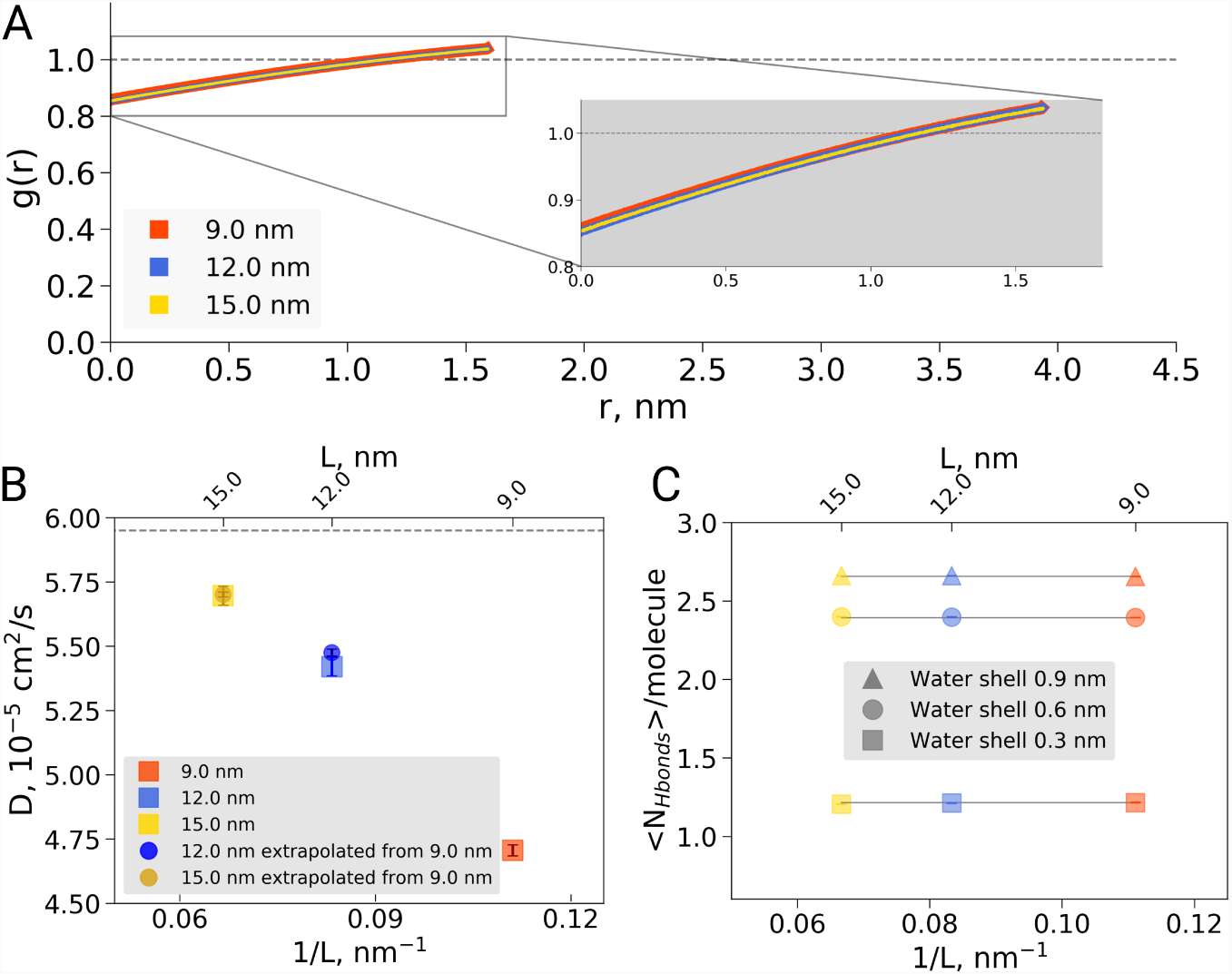
Radial distribution function (RDF), water diffusion and hydrogen bonds in hemoglobin simulations. (A) RDF for water oxygen atoms was calculated for the simulations in the boxes of different sizes. An equivalent volume for estimating the RDFs was used for every box size by cutting out a cubic box with an edge of 9 nm around the protein. (B) The water diffusion coefficient is plotted as a function of the box edge length. The square symbols mark the results obtained directly from simulations. The circles denote diffusion coefficients calculated from the value obtained from the 9 nm box simulations by means of extrapolation by adding a corresponding number of bulk water molecules (see main text). (C) Average number of water-water hydrogen bonds (cutoff value for the donor-acceptor distance of 0.33 nm) for three solvation shells around the protein: 0.3, 0.6 and 0.9 nm.

### Kinetics as a function of box size

To investigate if nevertheless the box size may affect transition kinetics, we have systematically investigated conformational transitions for a variety of additional systems including alternative setups of deoxy hemoglobin, each based on multiple independent trajectories.

We carried out MD simulations of deoxy hemoglobin in the Charmm36m force field for 4 different box sizes ranging from 8.25 nm to 16.3 nm for two different sets of protonation states for the histidine residues, starting from the T state (PDB entry 2dn2). In addition, simulations based on the El Hage et al. setup were carried out. Ten replicas of each state were simulated for statistics, except for the El Hage et al. setup with box sizes of 9 and 15 nm for which we did 20 replicas. Spontaneous transitions towards the R state were observed under each of the simulated conditions (Fig. 3A). Histidine protonation states as used by Hub et al. (4) were found to lead to generally faster transitions than histidine protonation as suggested by Kovalevsky et al. (5), in line with earlier simulations using the GROMOS96 43A1 force field (3, 6). Interestingly, the setup of El Hage et al. leads to generally slower transitions than either of these two protonation states. In contrast, no dependence on the simulation box size was observed for the transition times for any of the tested setups (Fig. 3).

**Figure 3:**
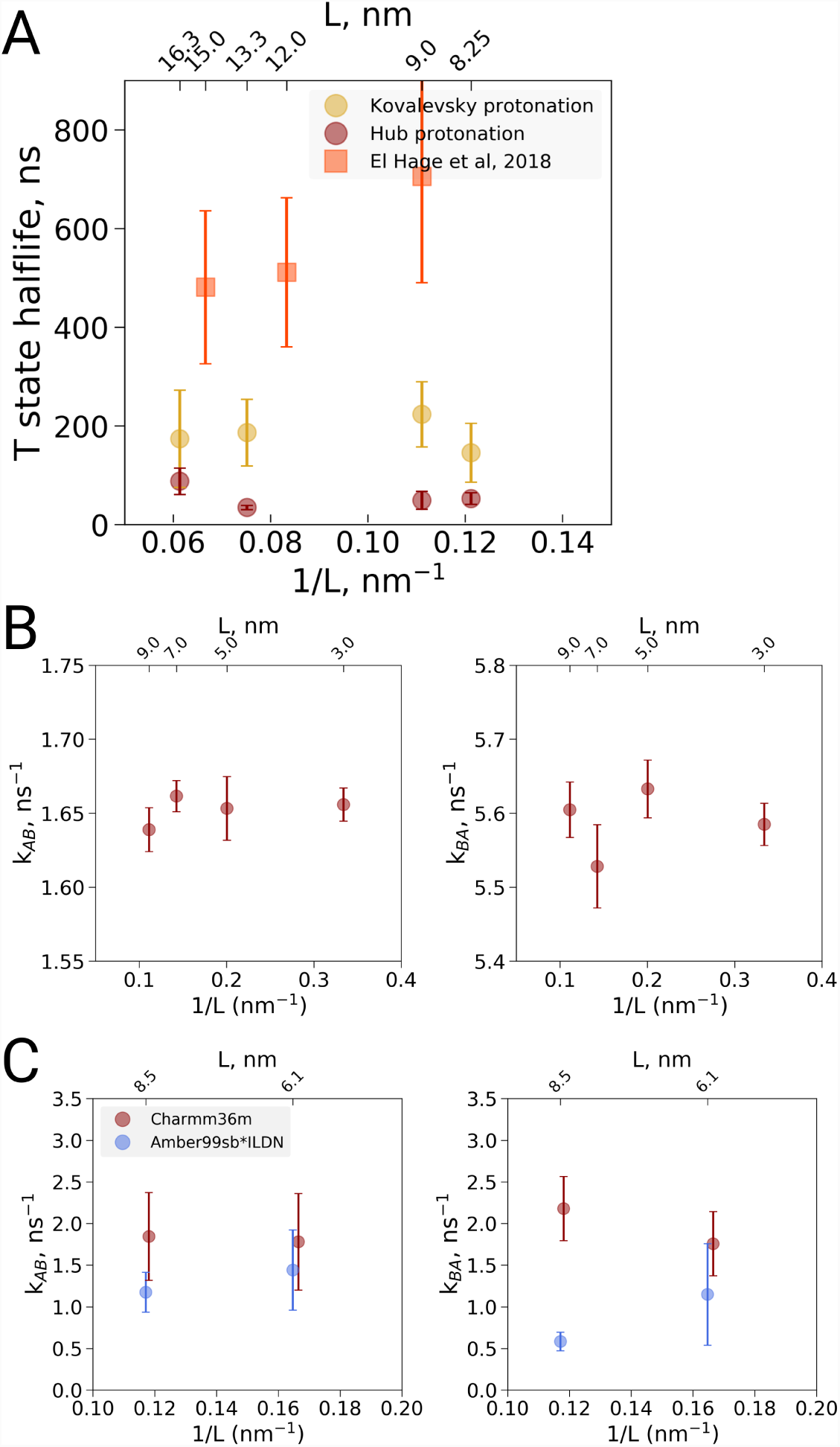
Kinetic analysis of hemoglobin, ubiquitin and alanine dipeptide. (A) Hemoglobin: average transition time from T to R, expressed as the time in which half of the replicas displayed the T to R transition, as a function of the simulation box size. Three different system setups were used for simulations. The Kovalevsky and Hub setups differ in the protonation of hemoglobin (3), both model the iron-proximal histidine interaction as a covalent bond. The El Hage setup is identical to the one used in (1), where the iron-histidine were not covalently bound. (B) Transition rates between the two minima in ubiquitin: rates from A to B (left) and from B to A (right). The reaction coordinate with the minima is depicted in Fig 4E,F. (C) Transition rates between the two minima in alanine dipeptide: rates from A to B (left) and from B to A (right). The reaction coordinate with the minima is depicted in Fig 4B,C.

### Factors influencing the T to R transition kinetics

In a preliminary investigation into the cause of the generally slower transitions in the El Hage et al. setup, we noted that the histidine protonation states were virtually identical to the Hub et al (4) setup and therefore are unlikely responsible for the slower transition times. The only difference in protonation state concerns His72, which is protonated on the delta position in both *α* subunits in the Hub setup, whereas it is protonated on the delta position on the *α*-1 subunit and on the epsilon position on the *α*-2 subunit in the El Hage et al. setup. Since *α*-His72 is exposed to the solvent, its protonation is unlikely to affect the T to R transition. Instead we identified two other differences that appear to slow down the transition. First, in the El Hage et al. setup an angular center of mass motion removal was employed on the protein, whereas in the Hub et al. and Vesper et al. a linear center of mass motion removal for the whole system was used. Second, whereas in the Hub et al. and Vesper et al. (3, 4) setup a covalent bond was modeled between the iron and the epsilon nitrogen of the proximal histidine for all four subunits, this interaction was modeled with non-bonded interactions in the El Hage et al. setup (1). Note that another difference in setup is the use of cubic and truncated dodecehedral boxes. Based on the current data, the box geometry does not appear to have a major effect, but more systematic analyses would be required for a definitive answer.

### Kinetics of conformational transitions in other systems

To further examine possible kinetic effects caused by the box size we in addition looked at two additional systems: alanine dipeptide and ubiquitin (Fig 4A-C). For alanine dipeptide, we counted transitions for the *ψ* dihedral, for ubiquitin transitions along the principal ‘pincer’ mode (7) were investigated. Also for these systems, no systemastic effect of the box size on the transition kinetics was observed.

**Figure 4:**
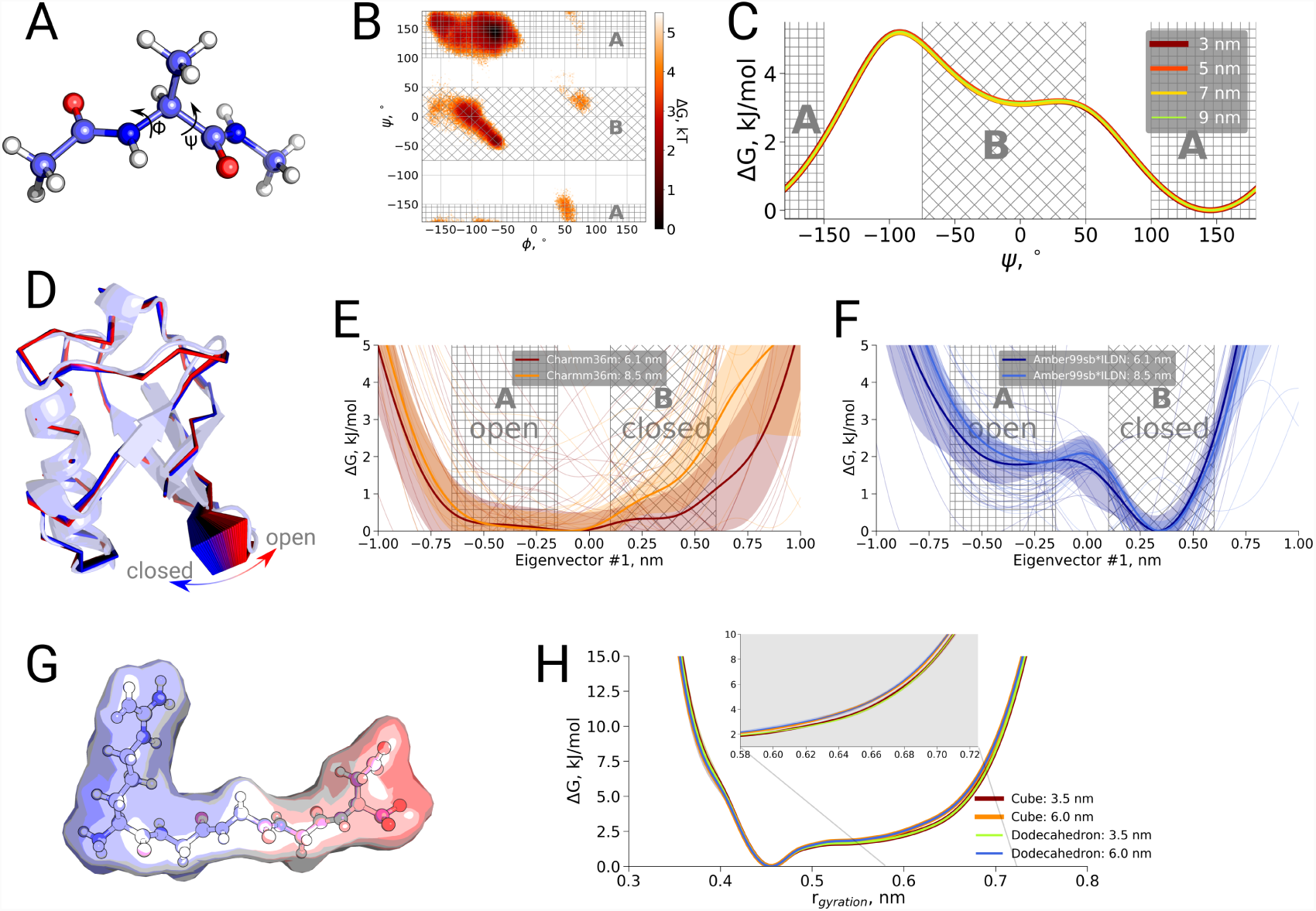
Thermodynamic analysis of hemoglobin, ubiquitin and RGGGD peptide. (A) Structure of the alanine dipeptide. (B) Free energy surface for alanine dipeptide simulated in the smallest box projected onto the backbone dihedrals marked in the panel (A). (C) Free energy profile along the *ψ* dihedral angle. (D) Structure of ubiquitin and an interpolation of the motion along the principal component with the largest variance (pincer mode). (E) Free energy profile projected on the pincer mode for ensembles generated with the Charmm36m force field. (F) Free energy profile for the Amber99sb*ILDN force field. (G) Structure of the RGGGD pentapeptide with the surface color corresponding to the surface charge: blue - positive, red - negative. (H) Free energy profiles for the pentapeptide projections as a function of the radius of gyration.

### Thermodynamics as a function of box size

Finally, we also investigated a possible box size on the thermodynamics of conformational transitions for alanine dipeptide, ubiquitin and the pentapeptide RGGGD. These systems have sufficiently small transition barriers to derive converged free energy profiles from unbiased simulations on the microsecond timescale. We constructed the pentapeptide RGGGD with the termini in their zwitterionic form, and thus carrying two positive charges on the N-terminus and two negative charges on the C-terminus, as a particularly sensitive test to long-range electrostatic effects on the conformational dynamics. In this case, no ions were added to the box, to avoid any screening effects. For the three systems, only minor effects, if any, of the box size on the obtained free energy profiles were observed (Fig 4).

## Conclusions

In conclusion, we have shown that the observation of El Hage et al. (1) that the stability of hemoglobin in the T state in the largest box size of 15 nm was not significantly different from the other investigated box sizes. In addition, a re-analysis of solvent hydrogen bonds, solvent radial distribution function and diffusion constants as a possible underlying hydrophobic or hydrodynamic effect showed that, when normalized to the analysis volume rather than the box volume, no significant box size effects were observed. Moreover, an analysis of the conformational transition kinetics in three hemoglobin setups as well as two additional systems showed no significant box size effects. Similarly, for three systems that allowed the investigation of thermodynamic effects, no significant box size effects were observed.

The results from hemoglobin simulations indicate that under the simulated conditions the R state is favored over the T state, with the histidine protonation states affecting the transition statistics, consistent with the Bohr effect (8) and previous simulations (3, 4).

Our analyses show that the dependence of the thermodynamic and kinetic properties on the box size is well within the level of statistical fluctuations for the observed statistics of 10-20 independent simulations at the microsecond timescale. The differences arising due to the different force fields, starting structures and chaotic nature of the simulations in this regime were found to be larger than the box size induced effects for the investigated systems.

## Materials and Methods

### Simulation setup

Three variants of the hemoglobin simulation setup were prepared: 1) El Hage (1), 2) Hub (4) and 3) Kovalevsky (5). The setups differed in the protonation state assignment and the protein-heme interaction treatment (see main text for details). The molecular dynamics simulations for the El Hage setup were performed in cubic boxes and used the same parameter set as reported in (1). For the Hub and Kovalevsky setups dodecahedral simulation boxes were used in combination with the molecular dynamics parameters compatible with the Charmm force field. The Charmm36m (9) force-field was used for all the simulations. Hemoglobin was solvated with TIP3P water, Na^+^ and Cl^-^ ions were added to neutralize the system and reach 150 mM salt concentration. Equations of motion were integrated with a 2 fs time step. The temperature was kept at 300 K by means of the velocity rescaling thermostat (10) with a time constant of 1.0 ps. All atoms in the system were assigned to one temperature coupling group. The Parrinello-Rahman barostat (11) was used to keep the pressure at 1 bar (time constant 5.0 ps). Long range electrostatics were treated by means of PME (12, 13) with a real space cutoff of 1.2 nm, a fourier spacing of 0.12 nm and interpolation order of 4. Van der waals interactions were cut off at 1.2 nm by smoothly switching off the force starting from 1.0 nm. Several noteworthy differences between the El Hage and other setup variants include a dispersion correction to energy and pressure which was used by El Hage et al, but was not employed for the other setups in our work. While El Hage setup used LINCS (14) to constrain all bonds, Hub and Kovalevsky setups constrained hydrogen atoms involving bonds only. The El Hage setup also removed angular center of mass motion of the protein during the simulation; this option was not used in other setups.

Simulations using the El Hage setup were performed in three cubic boxes with a cube edge of 9.0, 12.0 and nm. For the size of 9.0 nm, 20 independent simulations of 1 *µ*,s were performed, while for the size of 12.0, we used 10 simulations of 1 *µ*,s. For the size of 15.0 nm, 5 simulations reached 1 *µ*,s, while 15 additional runs reached 450-750 ns. The Hub and Kovalevsky setup runs were performed in dodecahedral boxes which have volume equivalent to the cubic boxes with an edge of 8.25, 9.0, 13.3 and 16.0 nm. For each box 10 simulations were performed reaching in total 2.8, 2.0, 0.9 and 1.2 *µ*,s for every box, respectively, in the case of Hub setup and 5.0, 4.0, 2.3 and 2.6 *µ*,s for every box in case of Kovalevsky setup.

The Alanine dipeptide, ubiquitin and RGGGD peptide simulations used the same simulation parameters as Hub and Kovalevsky hemoglobin setups. Alanine dipeptide was simulated in four cubic boxes with edges of 3.0, 5.0, 7.0 and 9.0 nm. For every box size 10 simulations 1 *µ*,s each were performed. Ubiquitin was simulation in two cubic boxes with edges of 6.1 and 8.5 nm. For ubiquitin 20 simulations 1 *µ*,s each were performed per box size: 10 runs starting from the 1ubq and 10 initialized with 1nbf structure. Furthermore, 20 additional ubiquitin simulations of 1 *µ*,s each were performed using Amber99sb*ILDN force field. RGGGD peptide was simulated in two cubic boxes with an edge length of 3.5 and 6.0 nm. For both box sizes 10 simulations of 1 *µ*,s were performed. In addition, the same simulation procedure for the RGGGD peptide was performed using dodecahedral boxes with the volume equivalent to the cubic boxes with an edge of 3.5 and 6.0 nm.

### Analysis

Root mean squared deviation (RMSD), radial distribution function (RDF), diffusion coefficient calculation and hydrogen bond counting were performed with the standard tools in the Gromacs (15) library. Prior to calculating RDF for the boxes with an edge of 12.0 and 15.0 nm, a cubic box with an edge of 9.0 nm centered at the protein was cut out from the original boxes. This ensured consistent normalization for all the box sizes. As for the hydrogen bond counting, the consistent normalization was ensured by considering only the waters around the protein in the shells at a distance of 0.3, 0.6 and 0.9 nm. For the box size dependence of the average diffusion constant, a bulk extrapolation based on the 9.0 nm box was used for the 12.0 and 15.0 nm boxes as described in the results section.

The error bars for hemoglobin halflife times, as well as alanine dipeptide and ubiquitin rate constants were calculated by means of bootstrap.

